# A Rationally Designed Aminoacyl-tRNA Synthetase for Genetically Encoded Fluorescent Amino Acids

**DOI:** 10.1101/065227

**Authors:** Ximena Steinberg, Jason Galpin, Gibran Nasir, Jose Sepulveda-Ugarte, Romina V. Sepúlveda, Fernando Gonzalez-Nilo, Leon D. Islas, Christopher A. Ahern, Sebastian Brauchi

## Abstract

The incorporation of non-canonical amino acids into proteins has emerged as a promising strategy to manipulate and study protein structure-function relationships with superior precision *in vitro* and *in vivo*. To date, fluorescent non-canonical amino acids (f-ncAA) have been successfully incorporated in proteins expressed in bacterial systems, *Xenopus* oocytes, and HEK-293T cells. Here, we describe the rational generation of an orthogonal aminoacyl-tRNA synthetase based on the *E. coli* tyrosine synthetase that is capable of encoding the f-ncAA tyr-coumarin in HEK-293T cells.

## INTRODUCTION

The emergence of chemical conjugation and genetic encoding techniques to label proteins with fluorescent probes has enabled significant advances in the mechanistic understanding of proteins in biochemical and cellular environments ^1^. Encoding large fluorescent proteins (e.g. GFP) as fusion protein products is experimentally straightforward, however, the relative size of the probes can alter the function and biology of the protein being studied. Alternatively, chemical conjugation of an expressed protein requires the labeling sites are solvent accessible and labeling of cytoplasmic sites often comes with significant background reactivity ^2^. A possible solution to these issues is the use of genetic code expansion to introduce a relatively compact fluorescent side chain as a non-canonical amino acid directly into the target protein in a site-specific fashion ^3^. Indeed, the rapid development of genetically encoded fluorophores as a non-canonical amino acids (ncAA) is emerging as a promising strategy to describe protein function under minimal perturbations in eukaryotic cells ^3,4^. The experimental strategy employs an orthogonal suppressor tRNA and an evolved aminoacyl tRNA synthetase (RS), often based upon the tyrosine pair, which can be used to encode the ncAA at virtually any site in the reading frame of the target gene. This pair is orthogonal to the translation system employed, meaning that an evolved TyrRs cannot acylate endogenous Tyr-tRNA molecules and the suppressor tRNAtyr is minimally acylated by host cell synthetases ^5^. The orthogonal tRNA has the appropriate anticodon to suppress the nonsense codon, thus allowing for an introduced amber codon of target genes in both prokaryote and eukaryotic systems ^3,5,6^. This approach has been successfully used for site-specific incorporation of f-ncAA into a soluble proteins in prokaryote cells (dansyl and hydroxycoumarin) ^4,7^, membrane proteins expressed in *Xenopus leavis* oocytes (bodipy and anap) ^8,9^, and proteins expressed in mammalian cells (dansyl and anap) ^10,11^. Hydroxycoumarin is notable because of its small size and high environmental sensitivity ^12,13^, however, no system yet exists for its incorporation via expanding genetic code in mammalian cells.

## RESULTS

The incorporation of a 7-hydroxycoumarin amino acid (TyrCoum) has been previously accomplished in *E.coli* using a *Methanococcus jannaschii* tyrosyl tRNA MjtRNA^Tyr^_CUA_ and a tyrosyl-tRNA synthetase (MjTyrRS) ^4^. To expand the possible applications for TyrCoum we sought to develop an tRNA/RS pair that is amenable to mammalian expression systems and thus started with the tyrosyl-tRNA synthetase from *E. coli* (EcTyrRS; PDBID 1X8X; Figure 1a) ^14^, in combination with a suitable tRNA from *B*. *stearothermophilus* (*Bs*tRNA_CUA_) ^6^. Evolved versions of this particular orthogonal pair allow for the efficient incorporation of a variety or aromatic ncAA, including p-benzoyl-L-phenylalanine (Bzp) and acetylphenylalanine (Azi) moieties in HEK-293T cells 15,16. Structural and structure-function information of the TyrRS was used as starting point for our *de-novo* design of a TyrCoum RS ^14,17^. First, comparable residues were identified within the amino acid binding region of the EcTyrRS (PDB 1X8X), MjTyrRS (PDB 1ZH6) ^18^ and structural models of the modified tyrosyl-tRNA synthetases MjCoumRS, EcAziRS, and EcBpaRS. These comparisons were useful as they informed the subsequent selection of RS mutations predicted to increase the size and flexibility of the catalytic domain. Four mutations are predicted to participate in substrate coordination (Y37G, L71H, D182G, and L186G) while five sites (L56E, T76G, S120Y, A121H, and F183I) are located near the catalytic domain and connector peptide region (Figure 1b). The Y37G mutation was introduced to destabilize tyrosine binding ^14,17^ and the additional D182G and L186G mutations served to further expand the binding pocket. The L71H and T76G are were reported in MjCoumRS (i.e. L65H and H70G) as crucial on the recognition of coumarin’s phenolic ring ^4^. S120Y and A212H, whose side chains are pointing out of the binding pocket, were introduced to promote flexibility within the binding region. The final engineered RS containing these nine mutations was termed EcCoumRS (Figure 1b). As a first pass, f-ncAA docking *in silico* to the EcCoumRS was examined by generating a molecular model using the structure of EcTyrRS as template. As expected according to the design strategy, the incorporated mutations produce a wider binding pocket with the potential to accommodate larger aromatic side-chains (Figure 1c and f). This structural analysis suggests that the f-ncAA substrate, TyrCoum, relies on a new set of hydrogen bonds to bind the catalytic site (Figure1 d and g) and that may preserve the orientation needed to form the end tRNA-amino acid complex (Figure 1e and h). *In silico* derived estimates of binding energy (ΔG_binding_=ΔG_apo_-ΔG_ligand-bound_), a value grossly related to the binding affinity^27^, suggests that the novel EcCoumRS could also be selective for TyrCoum (ΔG_binding_=-10 kCal/mol) over natural aromatic side chains (ΔG_binding_= -8.6, -8.3, -8.0 kCal/mol for Trp, Phe, and Tyr respectively). To test whether this new *Ec*CoumRS/*Bs*tRNA^tag^ pair was able to selectively incorporate TyrCoum western blot experiments were performed on HEK-293T cells expressing a mutant fluorescent protein (GFP^40^_TAG_) containing a mutation (Y40TAG) that precedes the protein cromophore ^16^. This strategy was used to test for rescue of the full-length fluorescent protein in the presence of the f-ncAA (500 nM; Figure 2a). Consistent with the *in silico* docking data, a second large fluorescent aromatic ncAA LysNBD (7- nitrobenz-2-oxa-1,3-diazol-4-yl) ^19^ was also encoded by the mutant RS but with lower efficiency (Figure 2a and b). Imaging data show that the suppressor pair rescues GFP^40^_TAG_ only in the presence of TyrCoum (Figure 2c and d), where a clear double staining is observed in cells eliciting coumarin signal (Figure 2e).

**Figure 1:**
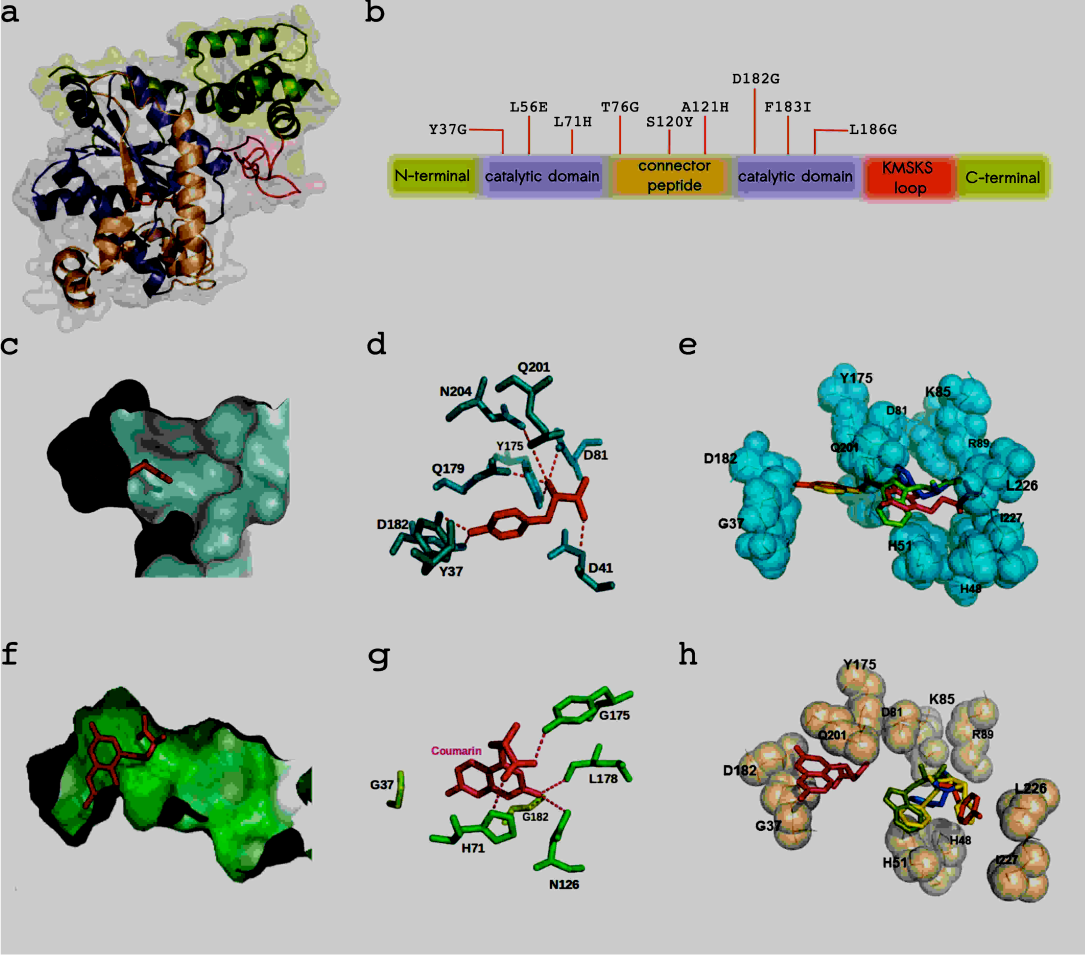
Figure 1: Homology modeling of the EcCoumRS enzyme and molecular docking studies. (a) Structure of the TyrRS enzyme (PDB: 1X8X) used to compute the CoumRS homology model. The Tyr ligand is shown in red. Color code depicts the different regions of the enzyme. C- and N-terminal regions in green, catalytic domains in blue, the linker between them in brown, and the KMSKS region in red. (b) Linear scheme of TyrRS highlighting the nine-engineered mutations present in the new enzyme. Color code is the same as in panel a. (c) Tyr ligand (red) in the binding pocket of TyrRS. (d) Network of amino acids stabilizing Tyr within the binding region. (e) Molecular docking showing that TyrRS binds Tyr leaving Phe, Trp, and Tyr-coumarin (yellow, green, and pink respectively) out of the principal binding region. (f) Tyr-coumarin ligand (pink) docked in the molecular model for CoumRS. The new enzyme presents a wider ligand-binding region when compared to the wildtype enzyme in c. (g) The network of amino acids stabilizing Tyr-coumarin is not well conserved when compared to the wildtype enzyme. Nevertheless, the lowest energy configuration suggests that the orientation of both ligands is similar. (h) View of the lowest energy configuration of Tyr/TyrCoum molecular docking shows that CoumRS binds Tyr-coumarin leaving the natural aromatic amino acids (Phe, yellow; Tyr, red; Trp, green) out of the principal binding region.

**Figure 2:**
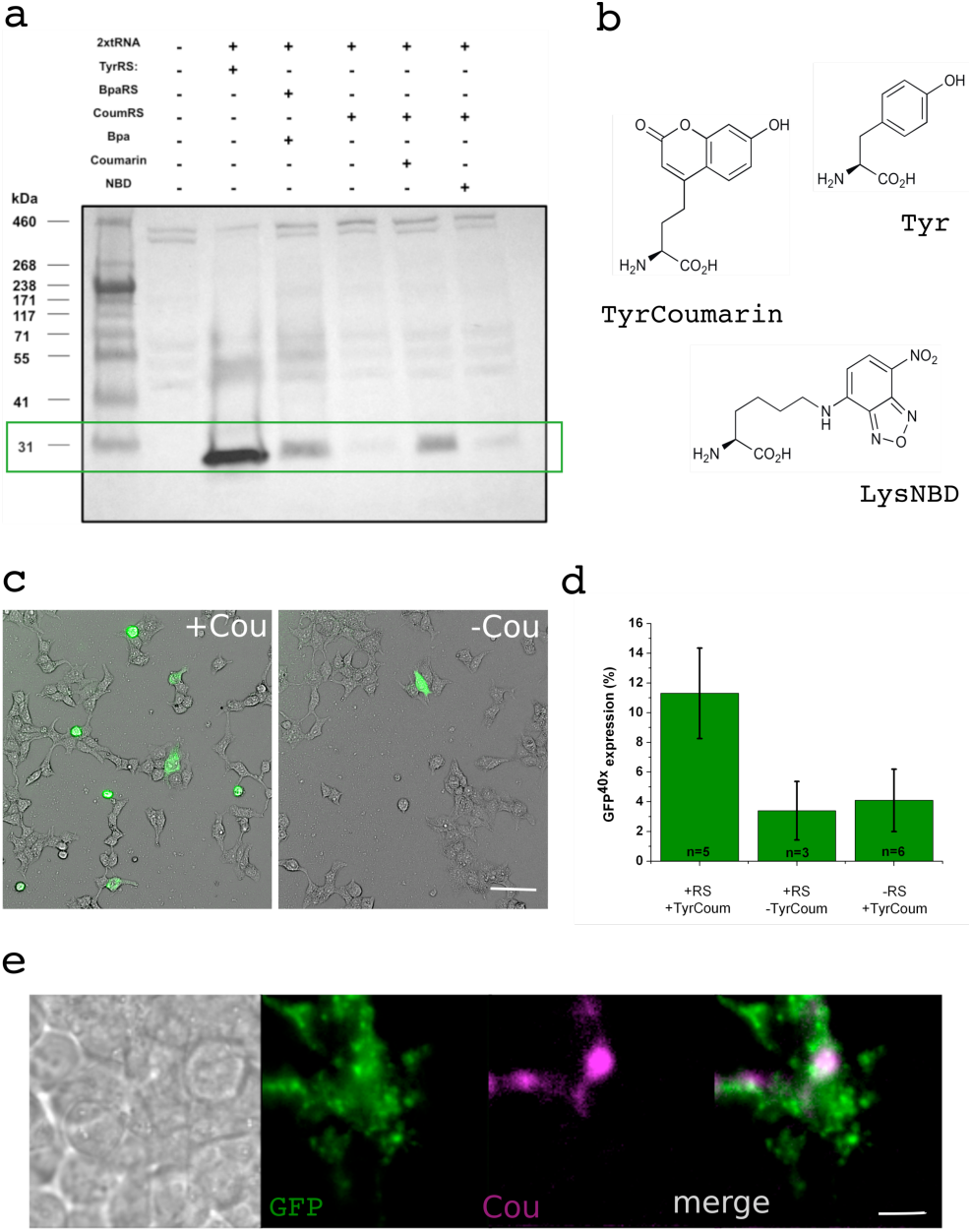
Amber codon suppression and incorporation of a coumarinyl aminoacid in HEK-293T cells. (a) Western blots for the detection of amber suppression in HEK-293T cells expressing the GFP^40^_TAG_ gene in the absence or presence of Tyr-coumarin and Lys-NBD (lower band at 30 kDa, green box). 2^nd^ lane is transfected with GFP^40^_TAG_ alone and the 3^rd^ lane represent the effect of expressing a TyrRS and the correspondant tRNA. The rescue of GFP^40^_TAG_ of the BpaRS (lane 4) is comparable to the rescue in the presence of CoumRS (lane 6). (b) Structures corresponding to the f-ncAAs used in this study and the natural amino acid tyrosine. (c) The suppression of the amber codon was estimated by analyzing the expression of GFP^40^_TAG_ on cultured HEK-293T cells. Images correspond to a representative filed of view in which transmitted light and fluorescent signals are merged for both the condition with (left) or without (right) Tyr-coumarin in the culture media. Bar = 40um (d) Quantification of GFP^40^_TAG_ expression in the absence or presence of Tyr-coumarin in the culture media or absence of CoumRS in the transfection mix. Bars correspond to SD. (e) Colocalization of coumarin and GFP signals. Low magnification images taken from transiently transfected HEK293T cells expressing GFP rescued with a coumarin side chain at position 40. The signal of coumarin fluorescence is absent in the cells not expressing GFP. Bar = 10um.

## DISCUSSION

The present study describes the design of an *in silico* designed amber suppressor tRNA/aminoacyl-tRNA synthetase pair for encoding Tyr-coumarin, *Ec*CoumRS. The approach takes advantage of the available structural and functional data on RS proteins from bacterial and eukaryotic systems that have been used previously to generate amber suppression-based genetic code expansion approaches. It has been reported that the substrate binding process in *Ec*TyrRS is supported by polar interactions where D182 binds the natural substrate (i.e. Tyrosine) and when the substrate is absent D182 is stabilized by polar interactions with neighbor residues ^17^. For the case of TyrCoum, the mutations on residues D182G, F183I and L186G not only widen the cavity hosting the bulkier substrate but also provide internal contacts in which G182 is limited to interact with L178 alone; I183 is at short distance from Q179, G180, and N187; and G186 can be stabilized by contacting Y190, G191 and V192. Our docking simulations suggest that G182 changes the orientation of the alpha helix containing it, favoring ligand binding. Deeper within the binding pocket, I183 and G186 are facing away from the substrate binding site, where the modeling suggests that they are coordinated by neighboring residues. Overall, the data support the feasibility of our design strategy of *de-novo* design where *Ec*CoumRS yields hydroxycoumarin-containing proteins for single cell imaging and biochemical scale expression and is able to encode the f-ncAA NBD, albeit with lower yields. Further experiments including *in vitro* evolution-guided selection of the *Ec*CoumRS might produce more selective enzymes for these two fluorophores. In summary, we are reporting a robust tool to incorporate an environmentally sensitive f-ncAAs such as hydroxycoumarin in eukaryotic proteins of living cells, useful will for live cell imaging and spectroscopic studies in native systems, without the need for additional purification procedures.

## EXPERIMENTAL PROCEDURES

### Molecular Biology

The gene encoding CoumRS was constructed with overlapping PCRs, using BpaRS as primary template. Oligonucleotides used for the aminoacyl-tRNA synthetase mutations were (mutated nucleotides are underlined):

a) 5’-CTTGTTCCATTG<u>GAA</u>TGCCTGAAACGC - 3’; (L56E).

b) 5’-AAGCCGGTTGCG<u>CAC</u>GTAGGCGGCGCG<u>GGG</u>GGTCTGATTGGC - 3’; (H71L, T76G).

c) 5’-TGTGGAGAAAAC<u>TATCAT</u>ATCGCGGCGAAC - 3’; (S120Y, T121H).

d) TTGCAGGGTTAT<u>GGCATC</u>GCCTGT<u>GGG</u>AACAAACAGT ACG - 3’; (D182G, M183I, A186G).

Oligonucleotides used for the incorporation of restriction sites into the cassette encoding the synthetase were (initiation and termination codons are underlined):

C-terminal

e) 5’-GTTTAAACTTAAGCTTGGTACCCCACC<u>ATG</u> -3’; (HindIII).

N-terminal

f) 5’-GACGACAAG<u>TAA</u>TCTAGAGGGCCCGTTTAA -3’; (XbaI).

The plasmid encoding the tRNA^UAG^ (pSVBpUC) was obtained from Thomas P. Sakmar Laboratory. PCRs were done with PfuUltra II Fusion HSD DNA polymerase from Agilent Technologies, according to the manufacturer’s instructions.

### Synthesis of f-ncAAs

Synthesis of TyrCoum. L-(7- hydroxycoumarin-4-yl) ethylglycine was obtained as described before ^4^. Synthesis of NBD-lysine. Na-(t-Butoxyccarbonyl)-L-lysine (308 mg, 1.25 mmol) was dissolved in a 3% solution of sodium bicarbonate in water (13 mL) in a 100 mL roundbottom flask equipped with a stir bar, and heated to 37°C. A solution of NBD-Cl (7- chloro-4-nitrobenz-2-oxa-1,3-diazole) (500 mg, 2.5 mmol) in 95% ethanol (35 mL) was slowly introduced via Pasteur pipet over 20 minutes. Following ∼8 hours of stirring the reaction mixture was acidified to pH ∼1 by the addition of 6M HCl and extracted 3 times with dichloromethane. The organic phase was washed with saturated sodium bicarbonate and the aqueous layer retained. 6M HCl was added with rapid stirring in the presence of dichloromethane to lower the pH until the desired product came out as a yellow cloud. The acid mixture was extracted with more dichloromethane, the organic fractions combined and dried over anhydrous magnesium sulfate and placed on a rotary evaporator until a brownish residue was obtained. The purity of the product was estimated at ∼90% based on TLC analysis. Without further purification the product was dissolved in dichloromethane (5 mL) and trifluoroacetic acid was slowly added (20 mL) and the solution was stirred at room temperature for 2 hours until TLC showed no t-BOC remained on the compound. The reaction was then concentrated under vacuum to yield a dark, oily solid.

### Bioinformatics

Structure models for AziRS, BpaRS and CoumRS were made with Modeller 9.13 (Andrej Sali Lab, UCSF), using as template the X-ray structure of E. coli TyrRS in complex with Tyr-AMS (PDBID 1X8X) ^20^. The sequence identity obtained from the pairwise alignment of E. coli TyrRS with AziRS, BpaRS, and CoumRS were 98.6%, 98.8%, and 95.8% respectively. Molecular docking simulations were performed in parallel using two different engines: i) Autodock Vina docking software (Molecular Graphics Lab, The Scripps Research Institute) and ii) the online server SwissDock (Swiss Institute of Bioinformatics, Molecular modeling group). Docking was guided within a grid calculated with Autodock Vina as: volume coordinates dimensions X=46, Y=24, Z=34, centered on X=12.825, Y=29.431, Z=19.000, with 0.436Å spacing. To first evaluate our docking strategy, we performed docking studies using Tyr ligand on the TyrRS crystal structure (1X8X, ligand removed) and calculate the RMSD for the position of the docked Tyr when compared to the Tyr at the crystal structure. To find a list of potential mutations implied in the ligand binding, an alanine screening was performed into Tyr binding pocket. The structures mutated were used as new receptors of Tyr and TyrCoum in docking calculations. From the obtained set of ΔG_binding_, we chose the best scores for docked ligands taking in consideration both the RMSD from Autodock Vina (RMSDd/l and RMSDu/b) together with the full-fitness value from Swiss-Dock. Molecular graphics and analyses of ligand structures were performed with the UCSF Chimera package (University of California, San Francisco).

### Cell culture

HEK293T cells, were cultured in DMEM (Gibco Inc.) supplied with 10% FBS (Gibco Inc.). Cells were prepared at (60-70)% confluence and transfected with lipofectamine 2000 following the instructions from the provider (Life technologies). The target gene was transfected 2-3 hours after an initial transfection of the tRNA^tag^/EcCoumRS pair. The f-ncAA was added to cell cultures to a final concentration of 0.5 to 5 μM together with the second transfection procedure.

### Western Blots

The existing media was aspirated and the cells washed with ice-cold PBS containing a cocktail of protease inhibitors (Roche), dispatched from the dish using a cell lifter and transferred to pre-chilled 1.5 mL tubes. The cells were then centrifuged (21000 × g; 4°C) for 5 minutes and the supernatant discarded. The cell pellets were vortexed in 100 μL of cell lysis buffer, and protease inhibitors for 10 min in a cold room. The samples were then centrifuged (21,000 × g; 4°C) for 20 min. Protein quantification was done using Bradford Protein Assay. 80 μg total protein of each sample (20 μg of TyrRS positive control) run on a 4-20% PAGE gel (pre-cast, Biorad). Proteins were then transferred onto a nitrocellulose membrane with cold pack (room temp); 5% milk blocking for 2 hr at room temp was followed by overnight incubation with the primary antibody (1:10,000; anti-GFP rabbit polyclonal; Synaptic Systems). Secondary incubation was performed for 2 hr at RT (1:10,000; Goat Anti-Rabbit IgG HRP Conjugate; Millipore). Membranes were incubated with house ECL for 2 minutes and exposed every 1 min for 20 min.

### Imaging

24 hours after target gene transfection, cells were disaggregated and plated on poly-L-lysine treated glass covers. The f-ncAA-containing media was removed at least 12h prior the experiment allowing cells to clear the soluble f-ncAA and the cells were imaged 36-48 hours after the second transfection using an inverted Olympus IX71 microscope. 405nm and 473nm solid-state lasers (Laser Glow) were focused to the backplane of a high-numerical aperture objective (Olympus 60X, N.A. 1.49, oil) by a combination of focusing lens. Fluorescence emission was collected by a EMCCD camera (Andor iXon^EM^ 860), after passing through an emission filter according to each acquisition wavelength band of interest (coumarin:450/70; YFP:540/40; Semrock). Excitation light was controlled by a 12mm mechanical shutter (Vincent associates). All imaging experiments were done at room temperature (20-22°C). Laser and focus control was performed using micromanager. Acquisition and digitalization was done with Andor Solis software. Percentage of suppression (i.e. GFP40x expression) was calculated by counting the number of green cells on 10 random fields of the sample and averaged.

## ACKNOWLEDGEMENTS

X. Steinberg is a MECESUP and CONICYT fellow. This work was supported by FONDECYT grants 1151430 (SB), 1131003 (FGN), Anillo Científico ACT-1107 (FGN) and ACT-1401 (SB), National Institutes of Health Grant GM106569 from NIGMS (CAA), PCCI12023, DRI-CONICYT/CONACYT (SB and LDI). CAA is a member of the Membrane Protein Structural Dynamics Consortium, which is funded by National Institutes of Health Grant GM087519 from NIGMS. LDI is supported by DGAPA-PAPIIT-UNAM grant IN209515. SB is part of CISNe-UACh and UACh Program for Cell Biology. The authors have no competing interests or other interests that might be perceived to influence the results and/or discussion reported in this paper.

## CONTRIBUTIONS

X.S., S.B, C.A.A and L.D.I. designed the project. X.S. performed experiments including biochemistry, molecular biology, imaging, and bioinformatics, together with data analysis. J.G. synthesize the ncAAs. G.N. performed WB. J.S-U., R.V.S., and F.G-N. performed bioinformatics. S.B. and X.S. analyze data. S.B. and C.A.A. provided reagents and materials. S.B, C.A.A, and L.D.I.

